# Real-time projections of Ebola outbreak size and duration with and without vaccine use in Équateur, Democratic Republic of Congo, as of May 27, 2018

**DOI:** 10.1101/331447

**Authors:** J. Daniel Kelly, Lee Worden, Rae Wannier, Nicole A. Hoff, Patrick Mukadi, Cyrus Sinai, Sarah Ackley, Xianyun Chen, Daozhou Gao, Bernice Selo, Mathais Mossoko, Emile Okitolonda-Wemakoy, Eugene T. Richardson, George W. Rutherford, Thomas M. Lietman, Jean Jacques Muyembe-Tamfum, Anne W. Rimoin, Travis C. Porco

## Abstract

**Background:** As of May 27, 2018, 54 cases of Ebola virus disease (EVD) were reported in Équateur Province, Democratic Republic of Congo. We used reported case counts and time series from prior outbreaks to estimate the current outbreak size and duration with and without vaccine use.

**Methods:** We modeled Ebola virus transmission using a stochastic branching process model with a negative binomial distribution, using both estimates of reproduction number *R* declining from supercritical to subcritical derived from past Ebola outbreaks, as well as a particle filtering method to generate a probabilistic projection of the future course of the outbreak conditioned on its reported trajectory to date; modeled using 0%, 44%, and 62% estimates of vaccination coverage. Additionally, we used the time series for 18 prior Ebola outbreaks from 1976 to 2016 to parameterize a regression model predicting the outbreak size from the number of observed cases from April 4 to May 27.

**Results:** With the stochastic transmission model, we projected a median outbreak size of 78 EVD cases (95% credible interval: 52, 125.4), 86 cases (95% credible interval: 53, 174.3), and 91 cases (95% credible interval: 52, 843.5), using 62%, 44%, and 0% estimates of vaccination coverage. With the regression model, we estimated a median size of 85.0 cases (95% prediction interval: 53.5, 216.6).

**Conclusions:** This outbreak has the potential to be the largest outbreak in DRC since 2007. Vaccines are projected to limit outbreak size and duration but are only part of prevention, control, and care strategies.

## Introduction

On May 8, 2018, the World Health Organization (WHO) announced the occurrence of an outbreak of Ebola virus disease (EVD) in the Democratic Republic of Congo (DRC).^1^ From April 4 through May 7, 21 suspected EVD cases were reported in Iboko and Bikoro, Équateur Province. On May 7, blood samples from five hospitalized patients had been sent to Kinshasa for Ebola-PCR testing, and two were confirmed PCR-positive.^2^ On May 21, vaccination of healthcare workers started.^3^ By May 27, the ring vaccination campaign was being rolled out, and there were 54 suspected, probable and confirmed EVD cases, including four confirmed cases in Mbandaka, the provincial capital of Equateur.^4^

This outbreak has several features that are worrisome for widespread transmission. Cases have been reported over a 168-kilometer distance, including the densely populated Mbandaka, a city of over 700,000 inhabitants situated on the Congo River and bordering Congo-Brazzaville.^5^ Moreover, travel to Kinshasa is frequent from Mbandaka. Given these risk factors, early epidemic growth profiles,^6^ and evidence of undocumented infection from previous outbreaks,^7,8^ the risk of a substantially larger outbreak cannot be ignored.

The factors causing epidemic growth to peak are debated. Delayed detection and resulting widespread distributions of EVD have significantly contributed to epidemic growth.^9^ Containment strategies which prioritize isolation over care can be counterproductive by discouraging sick individuals from presenting to high-mortality Ebola units.^10–12^ Unreported cases occurred at higher rates during the early period of the 2013-2016 West Africa outbreak,^13^ and urban transmission may also occur at a higher rate than rural transmission.^14^ Change to subcritical transmission (reproduction number below 1) tends to occur when Ebola response organizations deploy control, prevention and care measures,^15,16^ communities adopt more protective behaviors,^17,18^ and/or transmission decreases in a social network.^19,20^ In the current Ebola outbreak in DRC, scientific advances with rapid diagnostics and vaccines from the West Africa outbreak has the potential to limit Ebola virus transmission as they are being deployed.^21–23^

We used reported case counts from the current outbreak and/or time series from 18 prior outbreaks to estimate the total outbreak size and duration with and without the use of vaccines. These projections are intended to help organizations anticipate and allocate sufficient resources to the 2018 Ebola outbreak in DRC.

## Methods

We used the following methods to generate projections: a stochastic branching process model,^24^ statistical regression based on prior outbreaks, and Gott’s Law.^25^ On May 27, 54 EVD cases were reported in three locations (Iboko, Bikoro, and Mbandaka) (Figure 1). Based on EVD situation reports from DRC, we assumed the ring vaccination campaign would start on May 26.

**Figure 1.**
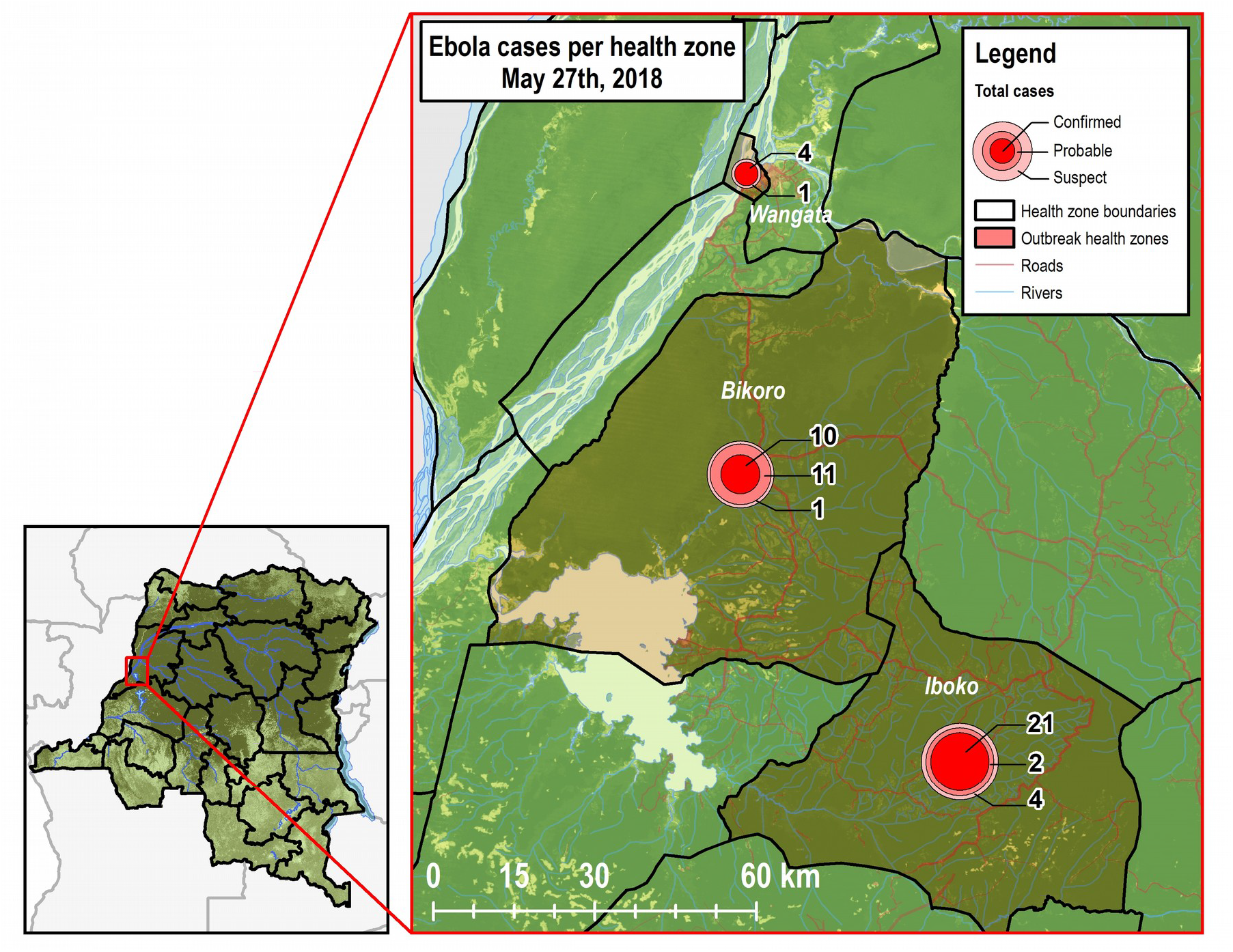
Map of cases of Ebola virus disease per health zone in Equateur Province, Democratic Republic of Congo

### Stochastic branching process model

We modeled Ebola virus transmission using a stochastic branching process model, parameterized by transmission rates estimated from the dynamics of prior Ebola outbreaks, and conditioned on agreement with reported case counts from the current outbreak to date. We incorporated high and low estimates of vaccination coverage into this model. Then we generated a set of probabilistic projections of the size and duration of simulated outbreaks in the current setting.

To estimate the reproduction number *R* as a function of the number of days from the beginning of the outbreak, we included reported cases by date from thirteen prior outbreaks and excluded the first historical outbreak reported in those countries (e.g., 1976 outbreak in Yambuko, DRC) (**Supplement 1**).^26–33^ As there is a difference in the Ebola response system as well as community sensitization to EVD following a country’s first outbreak, we employed this inclusion criterion to reflect the Ebola response system in DRC during what is now its ninth outbreak. We used the Wallinga-Teunis technique to estimate *R* for each case and therefore for each reporting date in the outbreak.^34^ The serial interval distribution used for this estimation was a gamma distribution with a mean of 14.5 days and a standard deviation of 5 days, with intervals rounded to the nearest whole number of days, consistent with the understanding that the serial interval of EVD cases ranges from 3 to 36 days with mean 14 to 15 days. We fit a curve *R* = *R_0_ e^−τd^* to each outbreak’s estimates of *R* by day *d*, initial reproduction number *R_0_* and quenching rate *τ* (**Supplement 2**).

We modeled transmission using a stochastic branching process model in which the number of secondary cases caused by any given primary case is drawn from a negative binomial distribution whose mean is the reproduction number R, and variance is controlled by a dispersion parameter *k*.^35,36^ All transmission events were assumed to be independent. The interval between date of detection of each primary case and that of each of its secondary cases is assumed gamma distributed with mean 14.5 days and standard deviation 5 days, rounded to the nearest whole number of days, as above. The pair of parameters *R_0_* and *τ* estimated for the different past outbreaks used, and dispersion parameter *k*, were used in all possible combinations (with *R_0_* and *τ* taken as a unit) to simulate outbreaks.

This model generated randomly varying simulated outbreaks with a range of case counts per day. The outbreak was assumed to begin on April 4 with a single case. The simulation process occurs as follows: proposed epidemic trajectories are generated in an initial step based on the above branching process, and these are then subsequently filtered by discarding all but those whose cumulative case counts match the known counts of the current DRC outbreak on known dates (Table 1). The filtration accepts epidemics within a range of 3 cases more or less than each recorded value. This one-step particle filtering technique produced an ensemble of model outbreaks, filtered on agreement with the recorded trajectory of the outbreak to date. This filtered ensemble is then used to generate projections of the eventual outcome of the outbreak.^14^

**Table 1.**

|  |  |
| --- | --- |
| May 7 (day 33) | 21 |
| May 11 (day 37) | 34 |
| May 14 (day 40) | 41 |
| May 18 (day 44) | 45 |
| May 20 (day 46) | 51 |
| May 23 (day 49) | 52 |
| May 27 (day 53) | 54 |

To model vaccination coverage with respect to total transmission (unreported and reported), we multiplied the estimate of vaccine effectiveness by low and high estimates of reported cases. In a ring vaccination study at the end of the West Africa outbreak, the overall estimated rVSV-vectored vaccine efficacy was 100% and vaccine effectiveness was 64.6% in protecting all contacts and contacts of contacts from EVD in the randomized clusters, including unvaccinated cluster members.^22^ We designed the stochastic model using vaccine effectiveness. Then, we used past estimates of the proportion of unreported cases to estimate the proportion of exposed individuals not covered by the vaccination process. Based on a Sierra Leonean study from the 2013-2016 outbreak,^13^ we estimated that the proportion of reported cases in DRC would rise over time from a low of 68% to a high of 96%. Given these low and high estimates of reported cases and the estimate of vaccine effectiveness, a low estimate of vaccination program coverage was 44% (68% * 64.6%) and a high estimate of vaccination program coverage was 62% (96% * 64.6%). We modeled the course of the outbreak with and without the vaccination program, with the program beginning on May 26, based on dates available from situation reports.^4^

In the 2013-2016 West Africa outbreak, the mean time from symptom onset to reporting was found to be approximately 4 days.^37^ Based on that estimate, we assumed a four-day period of transmission, with mean time of a transmission event falling two days before reporting. Based on this, we modeled transmission from a primary case to a secondary case using the value of *R* corresponding to two days before the reporting date of the primary case.

From 122,683,392 simulated outbreaks, 904 were retained after filtering on approximate agreement with DRC case counts. The simulated outbreaks that were retained after filtering were continued until they generated no further cases. We used this ensemble to derive a distribution of outbreak sizes and durations. We calculated mean and median values and 95% credible intervals using the 2.5 and 97.5 percentiles of simulated outbreak size and duration. We conducted the analyses using R 3.4.2 (R Foundation for Statistical Computing, Vienna, Austria).

### Regression model

For contrast with the stochastic model above, we conducted a simple regression forecast based solely on outbreaks of size 10 or greater. Time series for all 18 such prior outbreaks were obtained, including seven prior ones from DRC, dating back to 1976 (**Supplement 1 & 3**).^4,26–33,38–44^ No exclusions were made; no attempt to model specific features of this outbreak was conducted. The regression model predicted the outbreak size based on values of the outbreak size at a specific earlier time. The beginning of each outbreak was not reliably characterized; therefore, all time series were aligned on the day they reached 10 cases. In the current outbreak, we observed cases over the period from April 4 to May 27 (day 0 to day 53). May 27 corresponded to day 34 since reaching 10 reported cases. For the prior 18 outbreaks, we used linear interpolation to obtain the number of cases on day 34 (after reaching 10 cases). To reduce the influence of outliers and high leverage points, and to improve linearity, we calculated the pseudologarithm transform f(x)=arcsinh(x/2), asymptotically logarithmic but well-behaved at 0. We used non-parametric Theil-Sen regression (R-package *mblm*) followed by calculation of the resulting prediction interval for a new observation.^45,46^ Finally, we reported the median and 95% central coverage intervals for the prediction distribution conditional on the value being no smaller than the observed value on day 34. Sensitivity analysis was conducted using ordinary least squares regression. We conducted these analyses using R 3.4.2 (R Foundation for Statistical Computing, Vienna, Austria).

### Gott’s Law

We applied Gott’s Law to estimate the outbreak size using data from early in the current outbreak period.^25^ With Gott’s Law, we assume we have no special knowledge of our position on the epidemic curve. If we assume a non-informative uniform prior for the portion of the epidemic that still remains, the resulting probability distribution for the remaining number of cases *y* is P(Y=y)= Y_0_/y^2^. We estimated the median outbreak size, along with the two-sided 95% confidence interval.

## Results

As of May 27, 2018, there were 54 EVD cases (Table 1, Figure 1). Bikoko had ten confirmed cases, 11 probable cases, and one suspected case. Iboko had 21 confirmed cases, two probable cases, and one suspected case. Mbandaka had four confirmed cases and one suspected cases. In addition to cases, 906 contacts have been recorded and are actively being monitored. Twenty-five (46%) of 54 EVD cases have died.^4^

In the absence of any vaccination program, the projected median outbreak size based on the stochastic model was 91 cases (mean 161.0; 95% credible interval: 52, 843.5). Median duration of projected outbreaks was 112 days (mean 126.9; 95% credible interval: 54, 256.5). Using a lower estimate of 44% vaccination coverage, the median outbreak size was 86 cases (mean 91.4; 95% credible interval: 53, 174.3) and median duration was 96 days (mean 100.8; 95% credible interval: 54, 174.1). Using a higher estimate of 62% vaccination coverage, the median size was 78 EVD cases (mean 82.3; 95% credible interval: 52, 125.4), and the median duration was 89 days (mean 91.5; 95% credible interval: 54, 136.4) (Figure 2).

**Figure 2.**
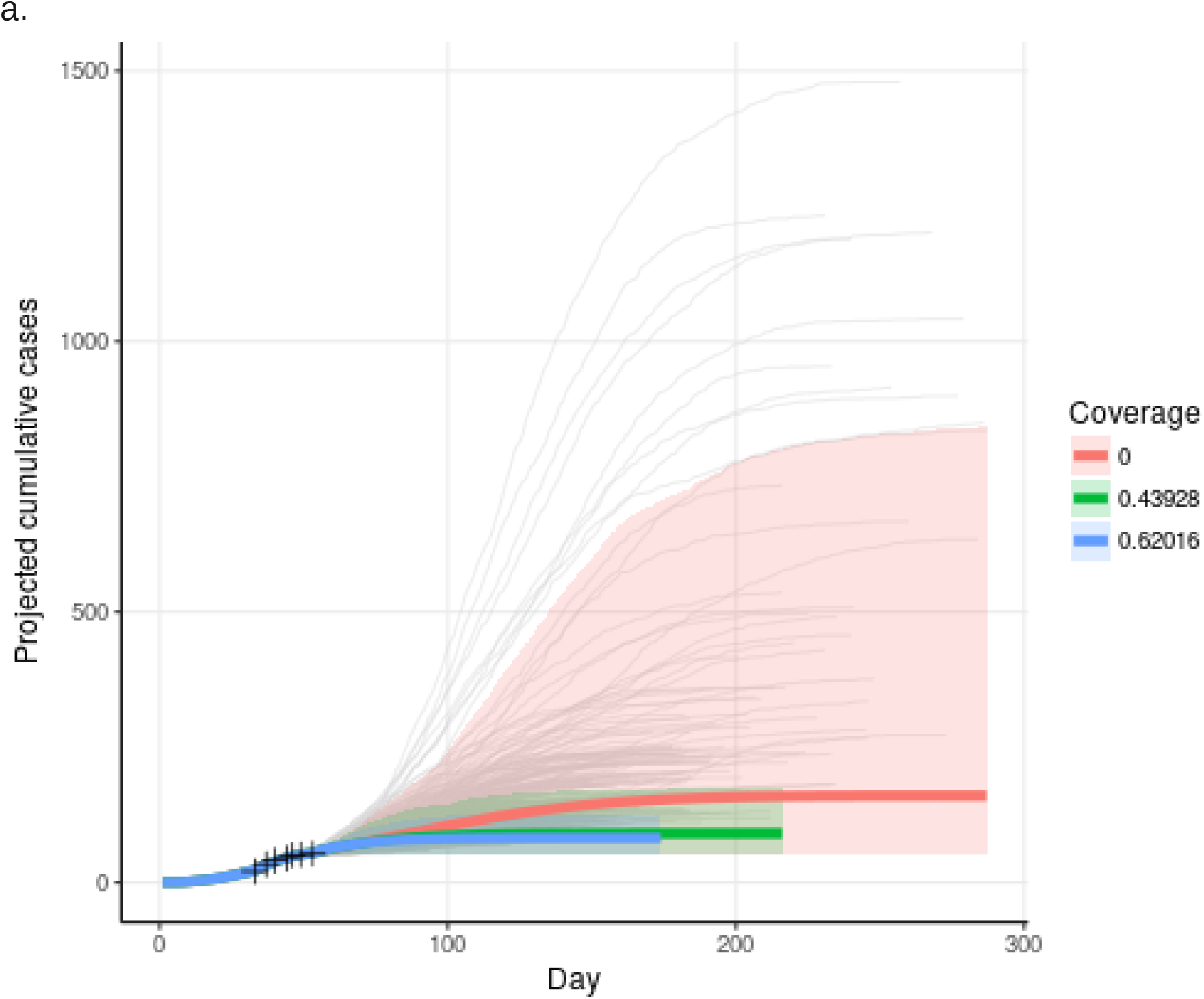
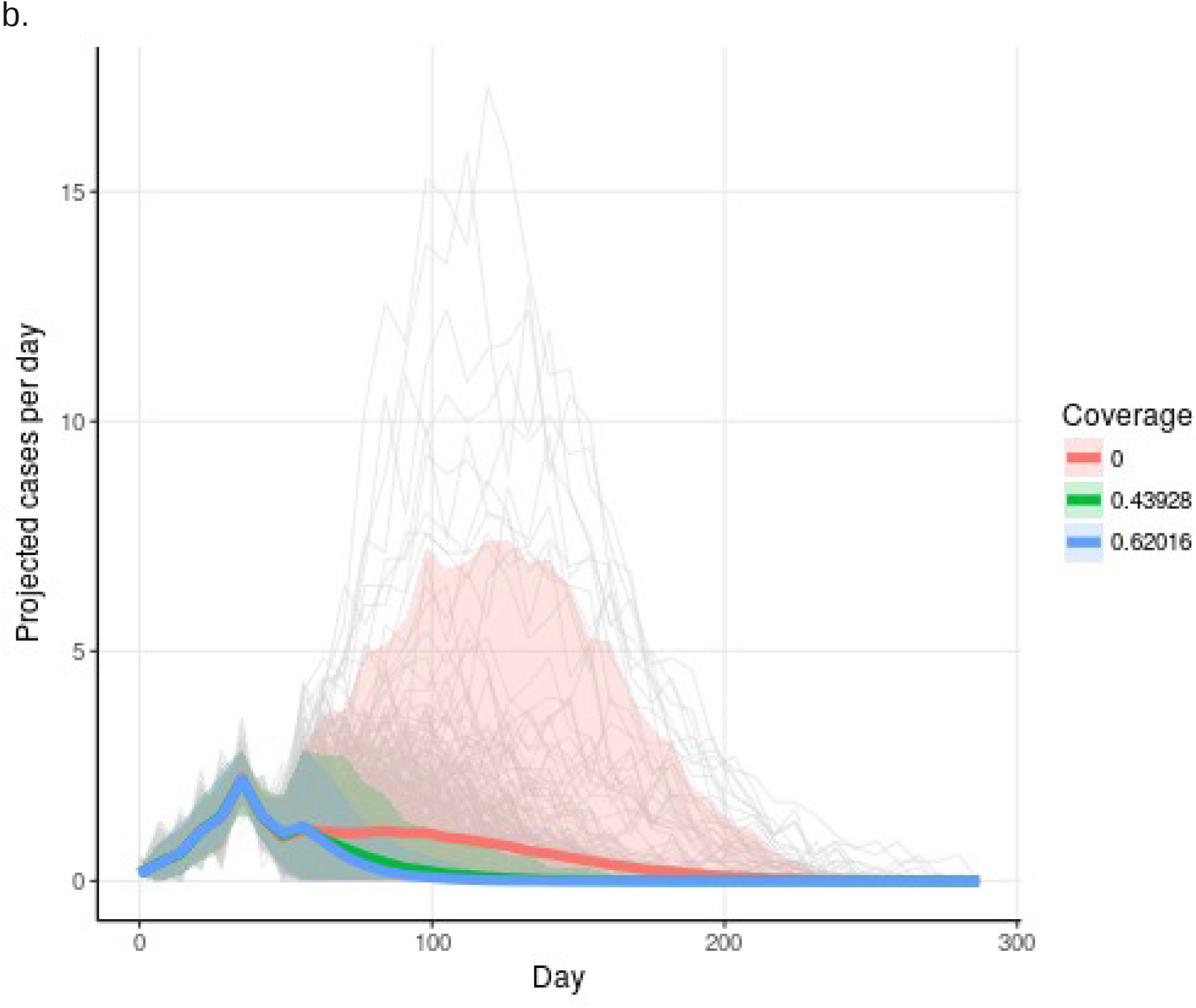
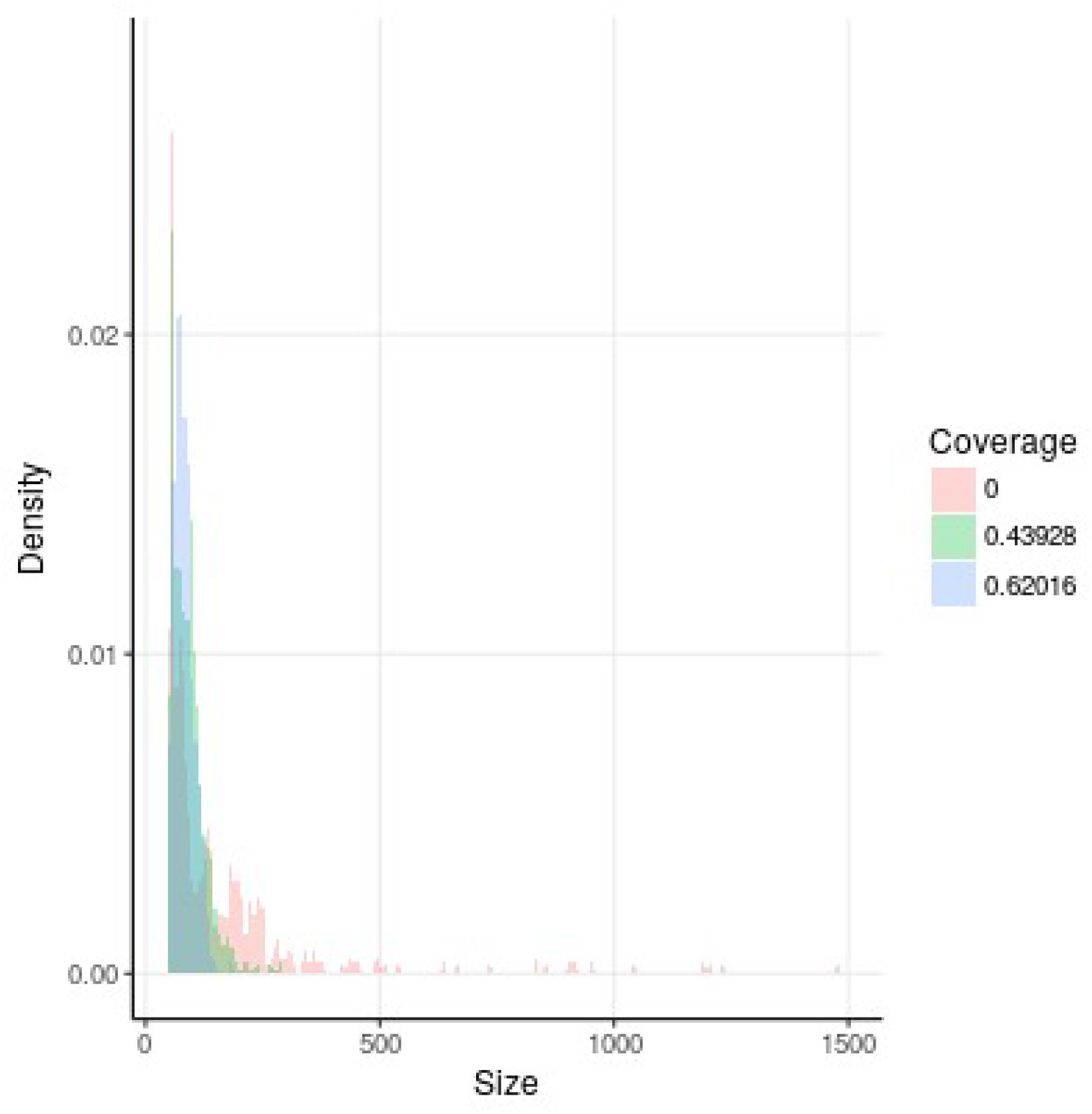
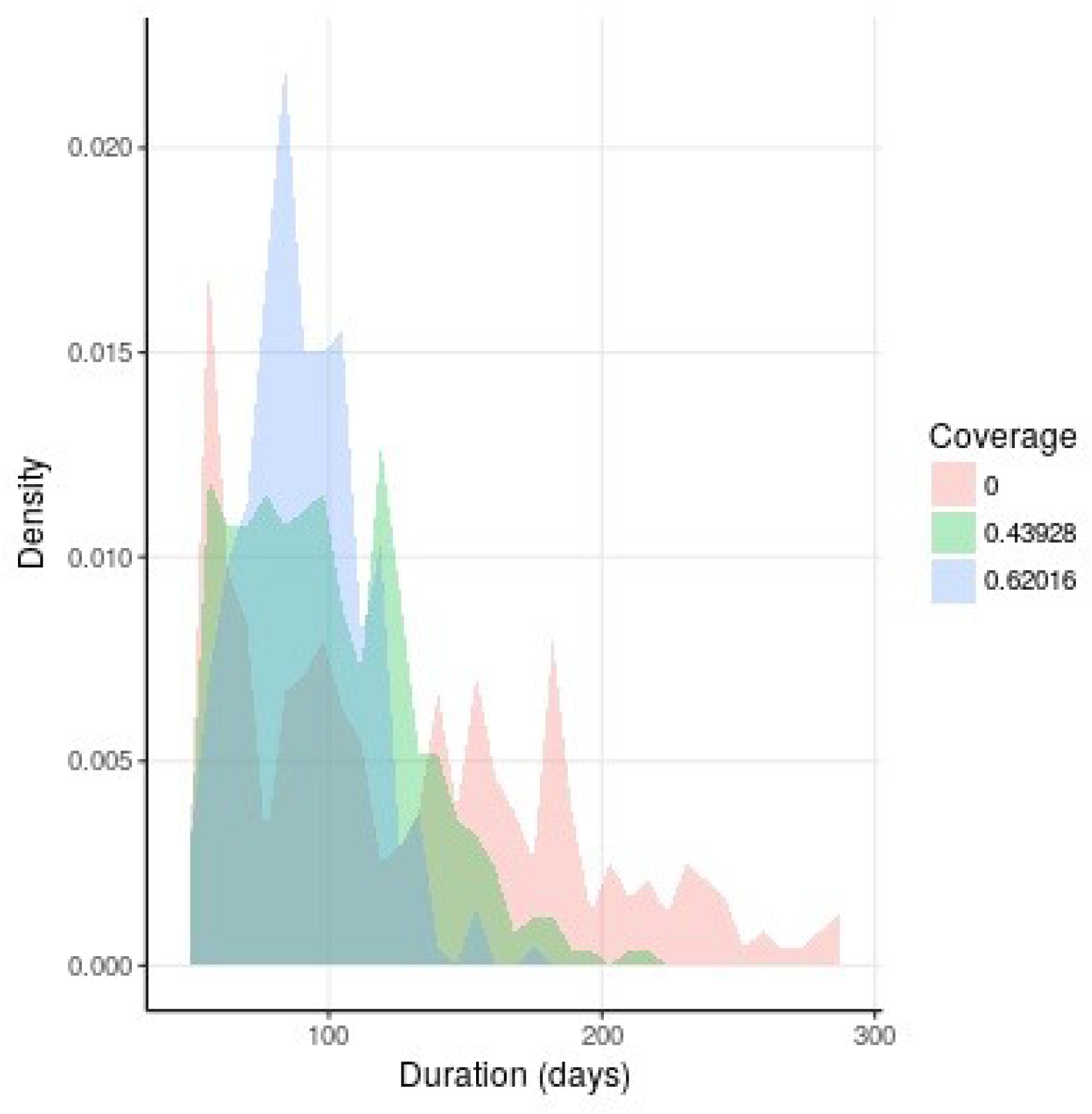
Distribution of outbreak projections from stochastic branching process model. **a**. Mean and credible interval of cumulative case count by day in model projections, by proportion of vaccine coverage. Target case numbers and dates are marked with + signs. **b**. Mean and credible interval of weekly average of per-day case count in model projections. **c**. Distribution of outbreak size over all model projections. **d**. Distribution of outbreak duration over all model projections.

With the regression based on past outbreaks, the median outbreak size was estimated to be 85.0 cases (95% prediction interval: 53.5, 215.6). Removal of data from the 2013-2016 West Africa outbreak from the regression yielded a median size of 71.4 cases (95% central coverage interval for outbreak size distribution: 53.0, 127.1), while use of ordinary least squares produced a median size of 114.4 cases (95% central coverage interval: 54, 433). One (5.6%) of the 18 prior outbreaks in our sample (the 2013-2016 West Africa outbreak) achieved a large size (over 28,000 cases).

Gott’s Law suggests that given 54 EVD cases, the median estimate of outbreak size was 108 EVD cases (95% CI: 55, 2160).

## Discussion

We estimated that the epidemic curve of this Ebola outbreak has peaked and new EVD cases in Équateur Province are declining at a rate consistent with subcritical transmission. This outbreak still has the potential to become the largest outbreak in DRC since 2007, regardless of vaccine use. Introduction of Ebola virus into Mbandaka with its over 700,000-person population resulted in multiple transmission chains and redoubles the importance of an adequate influx of support for response team efforts to implement interventions that control and prevent the spread of EVD. While real-time modeling projections historically overestimate outbreak size and duration,^47,48^ this study exemplifies how simple mathematical models can be useful for advising real-time decision making because they provide rapid projections and similar estimates of *R* as compared to complex models.^49^

Vaccine use, regardless of 62% or 44% coverage levels, is projected to provide a limited preventive benefit, which may be partially due to timing of implementation (after predicted peak of the epidemic curve). As such, the massive mobilization of resources to roll it out should not detract from the prioritization of patient and survivor care, lest we recapitulate the control-over-care paradigm that marred the response to the 2013-2016 outbreak in West Africa.^50–53^ Thus far, there has been a strong local and international response, and deployment of vaccines and rapid diagnostic tests (RDT) occurred early in response efforts.^4^ RDTs are being used to screen Ebola suspects while the vaccines are being administered to high-risk groups for EVD, including healthcare workers, contacts, and contacts of contacts. To limit epidemic growth from unreported cases, particularly those who have non-specific symptoms but are screened negative by the WHO case definition, more decentralized use of RDTs should be considered. We found that the impact of vaccines on curbing transmission would have been greater if vaccines were implemented prior to the peak of the epidemic curve (data not shown). Our assumptions about vaccine coverage in the current outbreak may be obviated if evaluations of vaccine use and coverage on transmission reduction are conducted.

There are limitations to our projections. Projection distributions are right-skewed, with long tails (and we therefore report the median instead of the mean). While there have been 22 observed EVD outbreaks with a case count greater than ten cases, we were unable to include all prior outbreaks in our estimates due to data availability.^44,54^ Note that the simple regression projection is based entirely on past outbreaks of Ebola virus disease (measured and reported in different ways), and cannot account for the improved control measures and vaccination in the way that a mechanistic model does. We included, however, as much real-time information into our estimates as possible, but situations such as the introduction of EVD into a large urban population and implementation of RDTs and vaccines are new to DRC. We did not include vaccination of healthcare workers in the stochastic model. Our estimates of vaccination effectiveness, reported cases, and mean time from symptom onset to reporting were obtained from West Africa not DRC studies. A strength of our approach was the use of multiple methods to estimate the outbreak size, even though Gott’s Law has not been validated for outbreak projections.

As of May 27, the epidemic curve of the EVD outbreak is thought to be declining, even though EVD transmission entered an urban setting. We do not believe this outbreak will be a repeat of the 2013-2016 West African one. Introduction of vaccines will limit transmission, even with lower levels of vaccination coverage, and DRC has the significant experience of eight prior outbreaks. As vaccine coverage is scaled up, an influx of support is warranted to support and bolster the evolving rapid response and be continued during the post-outbreak period. Stronger surveillance, preparedness, and health systems are needed to detect EVD cases before transmission threatens an even larger populace during future outbreaks.

## Acknowledgments

We thank the Ebola responders for their efforts in the current outbreak. We appreciate the Faucett Catalyst Fund for funding Ebola response efforts of the UCLA-DRC research program. TCP and LW are supported by the National Institutes of General Medical Sciences (NIGMS) under Award Number U01-GM087728.

## Authors’ contributions

J.D.K. contributed to concept and design, analysis, interpretation, drafting, revising, and final approval; L.W. contributed to concept and design, analysis, interpretation, drafting, revising, and final approval; R.W. contributed to acquisition, analysis, interpretation, revising, and final approval; N.A.H. contributed to interpretation, revising, and final approval; P.M. contributed to interpretation, revising, and final approval; C.S. contributed to acquisition, revising, and final approval; S.A. contributed to interpretation, revising, and final approval; X.C. contributed to acquisition, revising, and final approval; D.G. contributed to acquisition, revising, and final approval; E.T.R. contributed to concept and design, interpretation, revising, and final approval; G.W.R. contributed to concept and design, interpretation, revising, and final approval; T.L. contributed to concept and design, analysis, interpretation, revising, and final approval; J.J.M. contributed to interpretation, revising, and final approval; A.W.R. contributed to concept and design, interpretation, revising, and final approval; and T.C.P. contributed to concept and design, analysis, interpretation, revising, and final approval.

## References

1. World Health Organization. Emergencies preparedness, response. Ebola virus disease -- Democratic Republic of the Congo. 10 May 2018. Available at: http://www.who.int/csr/don/10-may-2018-ebola-drc/en/.

2. World Health Organization. Emergencies preparedness, response. 14 May 2018. Available at: http://www.who.int/csr/don/14-may-2018-eboladrc/en/.

3. Branswell H. Excitement over use of Ebola vaccine in outbreak tempered by real-world challenges. Stat. Health. 23 May 2018. Available at: https://www.statnews.com/2018/05/23/ebola-vaccine-drc-real-world-challenges/.

4. World Health Organization Regional Office for Africa. Health topics: Ebola virus disease. Available at: http://www.afro.who.int/health-topics/ebola-virus-disease.

5. Wikipedia. Mbandaka. 22 May 2018. Available at: https://en.wikipedia.org/wiki/Mbandaka.

6. Chowell G, Sattenspiel L, Bansal S, Viboud C. Mathematical models to characterize early epidemic growth: A review. Phys Life Rev 2016;18:66–97.

7. Richardson ET, Kelly JD, Barrie MB, et al. Minimally symptomatic infection in an Ebola ‘hotspot’: a cross-sectional serosurvey. PLoS Negl Trop Dis 2016;10:e0005087.

8. Kelly JD, Barrie MB, Mesman AW, et al. Anatomy of a hotspot: chain and seroepidemiology of Ebola virus transmission, Sukudu, Sierra Leone, 2015-16. J Infect Dis 2018.

9. WHO Ebola Response Team. After Ebola in West Africa--Unpredictable Risks, Preventable Epidemics. N Engl J Med 2016;375:587–96.

10. Richardson ET, Barrie MB, Nutt CT, et al. The Ebola suspect’s dilemma. Lancet Glob Health 2017;5:e254–56.

11. Chertow DS, Kleine C, Edwards JK, Scaini R, Giuliani R, Sprecher A. Ebola virus disease in West Africa--clinical manifestations and management. N Engl J Med 2014;371:2054–7.

12. Richardson ET, Barrie MB, Kelly JD, Dibba Y, Koedoyoma S, Farmer PE. Biosocial Approaches to the 2013-2016 Ebola Pandemic. Health Hum Rights 2016;18:115–28.

13. Dalziel BD, Lau MSY, Tiffany A, et al. Unreported cases in the 2014-2016 Ebola epidemic: Spatiotemporal variation, and implications for estimating transmission. PLoS Negl Trop Dis 2018;12:e0006161.

14. Krauer F, Gsteiger S, Low N, Hansen CH, Althaus CL. Heterogeneity in District-Level Transmission of Ebola Virus Disease during the 2013-2015 Epidemic in West Africa. PLoS Negl Trop Dis 2016;10:e0004867.

15. Funk S, Ciglenecki I, Tiffany A, et al. The impact of control strategies and behavioural changes on the elimination of Ebola from Lofa County, Liberia. Philos Trans R Soc Lond B Biol Sci 2017;372.

16. Lewnard JA, Ndeffo Mbah ML, Alfaro-Murillo JA, et al. Dynamics and control of Ebola virus transmission in Montserrado, Liberia: a mathematical modelling analysis. Lancet Infect Dis 2014;14:1189–95.

17. Winters M, Jalloh MF, Sengeh P, et al. Risk Communication and Ebola-Specific Knowledge and Behavior during 2014-2015 Outbreak, Sierra Leone. Emerg Infect Dis 2018;24:336–44.

18. Figueroa ME. A Theory-Based Socioecological Model of Communication and Behavior for the Containment of the Ebola Epidemic in Liberia. J Health Commun 2017;22:5–9.

19. Kiskowski M, Chowell G. Modeling household and community transmission of Ebola virus disease: Epidemic growth, spatial dynamics and insights for epidemic control. Virulence 2016;7:163–73.

20. Bellan SE, Pulliam JR, Dushoff J, Meyers LA. Ebola control: effect of asymptomatic infection and acquired immunity. Lancet 2014;384:1499–500.

21. FDA. OraQuick Rapid Antigen Test for Ebola. For Use Under Emergency Use Authorization (EUA) Only. Available at: http://www.fda.gov/downloads/MedicalDevices/Safety/EmergencySituations/UCM456912.pdf.

22. Henao-Restrepo AM, Camacho A, Longini IM, et al. Efficacy and effectiveness of an rVSV-vectored vaccine in preventing Ebola virus disease: final results from the Guinea ring vaccination, open-label, cluster-randomised trial (Ebola Ça Suffit!). Lancet 2017;389:505–18.

23. Dhillon RS, Srikrishna D, Garry RF, Chowell G. Ebola control: rapid diagnostic testing. Lancet Infect Dis 2015;15:147–8.

24. Galton F, Watson HW. On the probability of the extinction of families. J Royal Anthropol Inst 1875. 4; 138–144.

25. Gott, JR. Implications of the Copernican principle for our future prospects. Nature 1993. 363; 315–319.

26. Ebola haemorrhagic fever in Sudan, 1976. Report of a WHO/International Study Team. Bull World Health Organ 1978;56:247–70.

27. Baron RC, McCormick JB, Zubeir OA. Ebola virus disease in southern Sudan: hospital dissemination and intrafamilial spread. Bull World Health Organ 1983;61:997–1003.

28. Georges AJ, Leroy EM, Renaut AA, et al. Ebola hemorrhagic fever outbreaks in Gabon, 1994-1997: epidemiologic and health control issues. J Infect Dis 1999;179 Suppl 1:S65–75.

29. Khan AS, Tshioko FK, Heymann DL, et al. The reemergence of Ebola hemorrhagic fever, Democratic Republic of the Congo, 1995. Commission de Lutte contre les Epidémies à Kikwit. J Infect Dis 1999;179 Suppl 1:S76–86.

30. Boumandouki P, Formenty P, Alain E, et al. Prise en charge des malades et des défunts lors de l’épidémie de fièvre hémorragique due au virus Ebola d’octobre à décembre 2003. 2005. Available at: https://www.researchgate.net/publication/280954258_Prise_en_charge_des_malades_et_des_defunts_lors_de_l%27epidemie_de_fievre_hemorragique_due_au_virus_Ebola_d%27octobre_a_decembre_2003.

31. Nkoghe D, Kone ML, Yada A, Leroy E. A limited outbreak of Ebola haemorrhagic fever in Etoumbi, Republic of Congo, 2005. Trans R Soc Trop Med Hyg 2011;105:466–72.

32. Maganga GD, Kapetshi J, Berthet N, et al. Ebola virus disease in the Democratic Republic of Congo. N Engl J Med 2014;371:2083–91.

33. MacNeil A, Farnon EC, Morgan OW, et al. Filovirus outbreak detection and surveillance: lessons from Bundibugyo. J Infect Dis 2011;204 Suppl 3:S761–7.

34. Wallinga J, Teunis P. Different epidemic curves for severe acute respiratory syndrome reveal similar impacts of control measures. Am J Epidemiol 2004;160:509–16.

35. Blumberg S, Lloyd-Smith JO. Comparing methods for estimating R0 from the size distribution of subcritical transmission chains. Epidemics 2013;5:131–45.

36. Lloyd-Smith JO, Schreiber SJ, Kopp PE, Getz WM. Superspreading and the effect of individual variation on disease emergence. Nature 2005;438:355–9.

37. Wong JY, Zhang W, Kargbo D, et al. Assessment of the severity of Ebola virus disease in Sierra Leone in 2014-2015. Epidemiol Infect 2016;144:1473–81.

38. Rosello A, Mossoko M, Flasche S, et al. Ebola virus disease in the Democratic Republic of the Congo, 1976-2014. Elife 2015;4.

39. Outbreak(s) of Ebola haemorrhagic fever, Congo and Gabon, October 2001-July 2002. Wkly Epidemiol Rec 2003;78:223–8.

40. Camacho A, Kucharski AJ, Funk S, Breman J, Piot P, Edmunds WJ. Potential for large outbreaks of Ebola virus disease. Epidemics 2014;9:70–8.

41. World Health Organization. Weekly epidemiological record. Abonnement annuel. 2005. (43)80;369–376. Available at: http://www.who.int/wer/2005/wer8043.pdf.

42. Okware SI. Three ebola outbreaks in Uganda 2000-2011. University of Bergen dissertation for PhD. 2015. Available at: http://bora.uib.no/handle/1956/9239.

43. Althaus CL, Low N, Musa EO, Shuaib F, Gsteiger S. Ebola virus disease outbreak in Nigeria: Transmission dynamics and rapid control. Epidemics 2015;11:80–4.

44. Centers for Disease Control and Prevention. Ebola (Ebola Virus Disease): History of Ebola Virus Disease: 2014-2016 Ebola Outbreak in West Africa: Case Counts. Available at: https://www.cdc.gov/vhf/ebola/history/2014-2016-outbreak/case-counts.html.

45. Sen PK. Estimates of the regression coefficient based on Kendall’s tau. Journal of the American Statistical Association, 1968. 63(324);1379–1389.

46. Theil H. A rank-invariant method of linear and polynomial regression analysis. I, II, III. Nederl Akad Wetensch, Proc, 1950. 53;386–392, 521–525, 1397–1412. MR0036489.

47. Meltzer MI, Atkins CY, Santibanez S, et al. Estimating the future number of cases in the Ebola epidemic--Liberia and Sierra Leone, 2014-2015. MMWR Suppl 2014;63:1–14.

48. Butler D. Models overestimate Ebola cases. Nature 2014;515:18.

49. Wong ZS, Bui CM, Chughtai AA, Macintyre CR. A systematic review of early modelling studies of Ebola virus disease in West Africa. Epidemiol Infect 2017;145:1069–94.

50. Kelly JD, Mukadi P, Dhillon RS. Beyond vaccines: improving survival rates in the DRC Ebola outbreak. Lancet 2018.

51. Kelly JD, Richardson ET, Drasher M, et al. Food Insecurity as a Risk Factor for Outcomes Related to Ebola Virus Disease in Kono District, Sierra Leone: A Cross-Sectional Study. Am J Trop Med Hyg 2018;98:1484–8.

52. Richardson ET, Fallah MP, Kelly JD, Barrie MB. The predicament of patients with suspected Ebola - Authors’ reply. Lancet Glob Health 2017;5:e660–e1.

53. Richardson ET, Kelly JD, Sesay O, et al. The symbolic violence of ‘outbreak’: A mixed methods, quasi-experimental impact evaluation of social protection on Ebola survivor wellbeing. Soc Sci Med 2017;195:77–82.

54. Centers for Disease Control and Prevention. Outbreaks Chronology: Ebola Hemorrhagic Fever [Internet]. 2014. Available from: http://www.cdc.gov/vhf/ebola/resources/outbreak-table.html.

